# Aneuploidy Spectrum Analysis as a Primer for Copy Number Profiling of Cancer Cells

**DOI:** 10.1101/674929

**Authors:** Ahmed Ibrahim Samir Khalil, Anupam Chattopadhyay, Amartya Sanyal

## Abstract

**Motivation:** Hyperploidy and segmental aneuploidy are hallmarks of cancer cells due to chromosome segregation errors and genomic instability. In such situations, accurate aneuploidy profiling of cancer data is critical for calibration of copy number (CN)-detection tools. Additionally, cancer cell populations suffer from different levels of clonal heterogeneity and aneuploidy alterations over time. The degree of heterogeneity adversely affects the segregation of the depth of coverage (DOC) signal into integral CN states. This, in turn, strongly influences the reliability of this data for ploidy profiling and copy number variation (CNV) analysis.

**Results:** We developed AStra framework for aneuploidy profiling of cancer data and assessing their suitability for copy number analysis without any prior knowledge of the input sequencing data. AStra estimates the best-fit aneuploidy profile as the spectrum with most genomic segments around integral CN states. We employ this spectrum to extract the CN-associated features such as the homogeneity score (HS), whole-genome ploidy level, and CN correction factor. The HS measures the percentage of genomic regions around CN states. It is used as a reliability assessment of sequencing data for downstream aneuploidy profiling and CNV analysis. We evaluated the accuracy of AStra using 31 low-coverage datasets from 20 cancer cell lines. AStra successfully identified the aneuploidy spectrum of complex cell lines with HS greater than 75%. Benchmarking against nQuire tool showed that AStra is superior in detecting the ploidy level using both low- and high-coverage data. Furthermore, AStra accurately estimated the ploidy of 26/27 strains of MCF7 (hyperploid) cell line which exhibit varied levels of aneuploidy spectrum and heterogeneity. Remarkably, we found that HS is strongly correlated with the doubling time of these strains.

**Availability and implementation:** AStra is an open source software implemented in Python and is available at https://github.com/AISKhalil/AStra

## 1. Introduction

Cancer cells are plagued with microscopic and submicroscopic numerical and structural alterations of chromosomes. Chromosome missegregation and instability manifests in complex genetic makeups of cancer cells. They can simultaneously acquire near-polyploidy state or aneuploidy at whole-chromosome or portions (segments) of chromosomes [1, 2]. As a result, cancer cells frequently have polyploidy or hyperploidy profiles with widespread segmental aneuploidies [3-5]. Analysis of the Mitelman database indicates 86% of solid tumors and 72% of blood cancers exhibit whole-chromosome gain or loss, while 26% and 6% of them are near-polyploid, respectively [6]. Also, profiles of 47 ovarian cancer cell lines from the CCLE database showed that up to 74% of their genomes are altered [7]. Furthermore, cancer cell populations are genetically highly heterogeneous. Sometimes they remain stable with aneuploid karyotype, and in other times, they undergo continuous genetic alterations over passaging time and culture conditions [8]. Same cell line cultured in different labs exhibits karyogram changes [9]. These facts suggest that cancer cells undergo an adaptive and evolutionary process even under controlled culture environment to generate clonal variability at copy number, gene expression and phenotypic levels [10]. Taken together, all these alterations give rise to ‘mosaic’ karyotypes of cancer cell populations.

Ploidy information is associated with cancer progression and their varied sensitivity to chemotherapy. Aneuploidy, in general, is considered as a marker of poor prognosis and their presence correlates with chemoresistance [11]. In fact, antiproliferative compounds targeting aneuploid cells serve as a new chemotherapeutic strategy for cancer cells [12]. Furthermore, characterization of the aneuploidy profile forms the basis of CNV detection for single-sample cancer data[13]. Most read depth (RD)-based tools use the global RD median for setting the CN reference (2N) [14, 15] or the ploidy states [16, 17]. However, the global median may not accurately approximate any of the ploidy states, especially in case of hyperploid cancer cells. Moreover, some of these tools require the information of CN gain/loss percentage for better modeling of the RD signal. Therefore, it is necessary to estimate the overwhelming extent of aneuploidy spectrum for calibrating the CNV detection tools.

The chromosomal aneuploidy and microscopic-level segmental gain or loss are classically analyzed by karyotyping. Deviation from ‘normal’ karyotype is a common hallmark of cancer cells [18, 19]. Whole-genome ploidy level with modal chromosome number has been routinely characterized for most cancer cell lines using imaging techniques by individual labs and cell repositories. However, cancer cells undergo clonal evolution over time and therefore ploidy characterization may not be stable and reliable. Moreover, karyotyping is a low-resolution technique. Therefore, standard karyotype cannot distinguish between two genomes with normal chromosome complement but harboring different submicroscopic chromosomal aneuploidies. Taken together, the dynamics of these microscopic and submicroscopic alterations represents timestamp ‘barcode’ of aneuploidy spectrum of cancer cells. A rough estimation of this spectrum can form the background knowledge needed for accurate downstream sequence analysis of cancer cell line.

The advent of array- and next-generation sequencing (NGS)- based techniques allowed higher-resolution estimation of the complete repertoire of aneuploidy at different structural levels. Most genome-level ploidy tools compute ploidy levels by assessing the distribution of allele frequency at biallelic single-nucleotide polymorphisms (SNPs) [20-22]. These SNP-based tools require very high-coverage sequencing data. As a result, they were successfully applied to small genomes like yeast but are less adaptable in human scenario, especially for low-coverage sequencing data. Additionally, PloidyNGS estimates ploidy by assuming different ratios of biallelic SNPs at different ploidy levels. However, this can bring ambiguity in defining the diploid versus tetraploid with 0.5/0.5 ratios. Recently, nQuire was developed to estimate ploidy by fitting the coverage signal to *pre-determined* frequency distributions with *fixed* contribution from each ploidy state [23]. Nevertheless, nQuire cannot capture the hyperploidy profiles which have *variable* combinations of different ploidy levels. Hence, it is necessary to capture the contribution of each ploidy level into the complete aneuploidy profile (aneuploidy spectrum) without any pre-defined assumption of whole-genome ploidy. Additionally, ploidy tools are implicitly built on a fundamental assumption that cancer cell lines are clonal in origin, and therefore, they are ‘homogeneous’ in nature. The pervasive presence of cellular variability and genetic heterogeneity in cancer cells [24] necessitate estimating the homogeneity level in cancer population as a condition for accurate ploidy estimation.

Here, we introduce AStra (**A**neuploidy **S**pectrum (detection) **t**hrough **r**ead depth **a**nalysis), a Python-based software for capturing the complete aneuploidy profile of cancer genome without an explicit assumption of the ploidy level of the input sequencing data. AStra is based on the fundamental reasoning that most genomic segments belong to integral CN states. AStra scans the RD signal to find the CN reference that best segregates genomic segments into distinct copy-number states to estimate the aneuploidy spectrum. We employ this spectrum to compute several CN-associated attributes. The homogeneity score measures the reliability of the input cancer data for downstream aneuploidy and CNV analysis. The tool also infers the whole-genome ploidy level, the CN correction factor and CN gain/loss percentage for calibrating CNV detection tools. To sum up, AStra provides the prerequisite knowledge of the copy number-associated features for the DOC-based analysis of cancer cells.

## 2. Methods

### 2.1 AStra framework

AStra utilizes the RD signal of the input data to define its aneuploidy spectrum and to compute its important features for accurate ploidy and CNV analysis. The main key for defining the aneuploidy spectrum is the accurate estimation of CN reference (2N) by scanning the RD signal. CN reference should be defined as the RD value that segregates RD signal distribution into segments of integral CN states. In our pipeline, NGS reads are used first to compute the RD signal. This RD signal is clustered using Pruned Exact Linear Time (PELT) method [25] into primary segments of different RD values. Then, we employ 10 different unimodal/multimodal distributions to define the candidate CN references and the associated CN states (Fig. 1a). Under each candidate reference, we merge the adjacent primary segments with same CN state into large contiguous segment to generate the aneuploidy profile. Then, we compute the Centralization Error (CE) of that aneuploidy profile as the weighted summation of differences between the estimated CN of segments and their CN states (Fig. 1b). The correct CN reference is the one that achieves the minimum CE. Once the CN reference is settled, we compute the ploidy spectrum as the percentages of each ploidy level into the complete aneuploidy profile. Finally, the homogeneity score (HS) is calculated as the percentage of genomic segments with estimated CN around their integral CN states (Fig. 1c). We also define the whole-genome ploidy (‘diploid,’ ‘triploid,’ or ‘tetraploid’) and determine the validity of usage of the global median (RD signal) as CN reference.

**Fig. 1.**
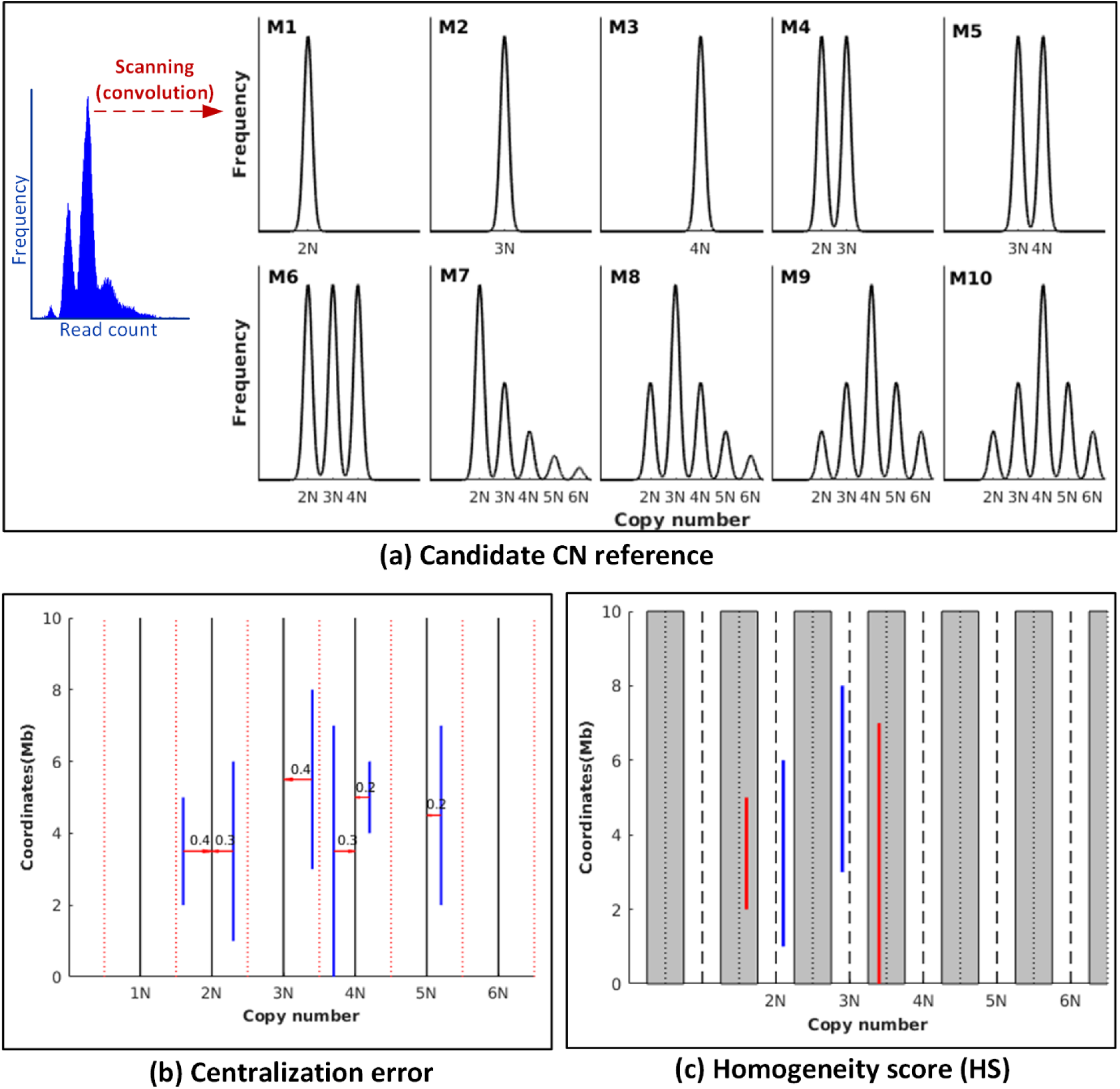
AStra framework. **(a)** Representative multimodal coverage distributions (M1 to M10) which are utilized to detect CN reference by scanning the input RD signal distribution. The RD frequency distribution of A427 cell line is shown as an input example. **(b)** Illustration to compute centralization error which allows to obtain the best-fit aneuploidy profile. Each blue line represents a genomic segment while the red arrow is the error weight of this segment (the difference between the CN of that segment and the nearest CN state). **(c)** Illustration to calculate the homogeneity score (HS). Each blue line represents a segment with estimated CN within 0.25N from the nearest CN state (homogeneous segment) whereas each red line represents a segment with estimated CN >0.25N from the nearest CN state (heterogeneous segment).

### 2.1.1 RD frequency distribution

Cancer cells exhibit multimodal distribution as a combination of the underlying unimodal distributions at different CN states. Therefore, we build 10 RD frequency distributions as weighted summation of Gaussian/Normal distributions centralized at different CN states to compute the candidate CN reference (Fig. 1a):

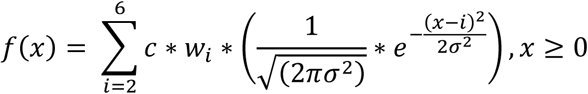

such as

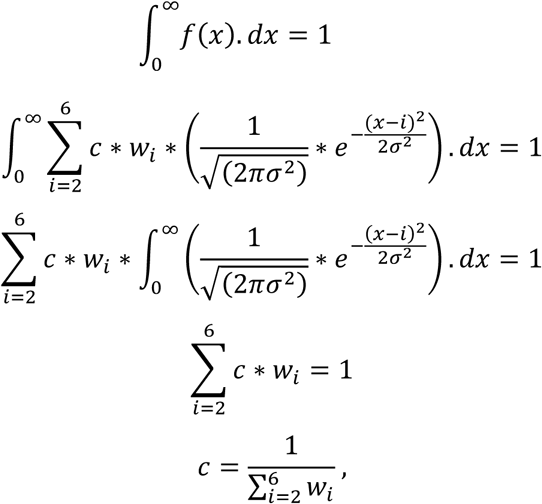

where *i* is the copy-number state (2, 3, 4, 5, 6), *w*_*i*_ is the weight of the Gaussian distribution at state *i* (0, 1, 0.5, 0.25, 0.125, 0.0625) and *c* is a constant for normalization of the probability distribution function. The standard deviation (σ) is 0.5/3 to make *f* (*x*) 0 at the boundaries of copy-number intervals (1.5, 2.5, 3.5, …).

### 2.1.2 Estimation of candidate CN reference

For each of the 10 RD frequency distributions, candidate CN reference is defined as the RD value that achieves the maximum correlation between the RD frequency distribution *r* (*x*) and the multimodal distribution *f* (*x*). The interval of RD values [s, e] is divided into *m* equally spaced RD values. At each RD value *k*, a multimodal distribution *f*_*k*_(*x*) is generated assuming this is the copy number reference (2N). And, a rank *R*_*k*_ is computed as the convolution between *r* (*x*) and *f*_*k*_(*x*):

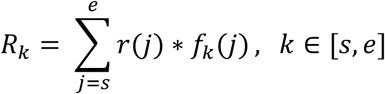

Finally, RD value *k* with the maximum rank *R*_k_ is chosen as the candidate CN reference.

### 2.1.3 Centralization error

Given the candidate CN reference, we merge the genomic segments to divide the genome into contiguous segments of distinct CN states to form the chromosomal aneuploidy profile. Under the correct CN reference, most of these segments should have CN around integral CN states. CE measures the degree of localization of these segments near CN states (Fig. 1b):

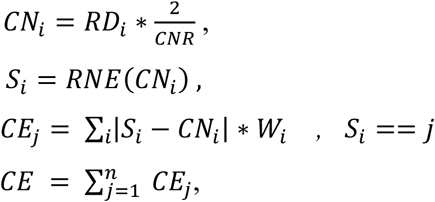

where *i* is the segment index, *j* is the CN state/interval, *RD*_*i*_ is the median RD per segment, *CN*_*i*_ is the copy number of the segment, CNR is the candidate CN reference, *S*_*i*_ is the CN state of the segment (round-to-nearest value of *CN*_*i*_), *W*_*i*_ is the width of the segment *i*, and *CE*_*j*_ is the centralization error of segments of CN state *j*.

### 2.1.4 Characterization of the aneuploidy spectrum

The aneuploidy profile, with minimum CE, is decoded into the aneuploidy spectrum which contains the percentage of genome segments in each CN state/interval. HS measures the degree of localization of genomic segments into integral CN states. HS is computed as the percentage of segments that are around the integral CN states (Fig. 1c):

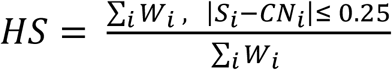

where *i* is the segment index, *CN*_*i*_ is the copy number of the segment, *S*_*i*_ is the CN state of the segment, and *W*_*i*_ is the width of the segment *i*. Sequencing data with high HS are suitable for CNV and ploidy analysis. Finally, we infer the ploidy level (‘diploid’, ‘triploid’, ‘tetraploid’) as the CN state harboring maximum percentage of genomic segments. Also, we compute the ‘CN correction factor’ as the RD signal ratio between the global median and the ploidy state. If any CNV tool computes CN reference solely based on the global median, this correction factor can be used to correct CN reference and copy number estimates of genomic segments.

## 3. Results

### 3.1 Coverage is not the only criteria for accurate aneuploidy profiling and copy number analysis of cancer cells

Cancer cells suffer from a degree of hyperploidy and segmental aneuploidy due to chromosomal instability. This results in a genome which contains a collection of genomic segments with different CN states. In such a situation, the copy number profiling can still be performed using appropriate modeling of RD signal by a multimodal distribution if and only if the CN profiles are well separated into integral CN states. However, this condition depends on the other source of biases in cancer sequencing data. The main bias originates from the purity of the sample, clonal variations, and heterogeneity of the cancer cells/samples which adversely affects peak separation. In addition, coverage signal of cancer cells may be affected by wave artifacts which may contribute to signal variations and peak separation [26]. In low coverage data, the effects of heterogeneity and wave artifacts are magnified due to the sparseness of reads representing each region of the genome. Consequently, it is a prerequisite to determine the suitability of a sequencing data for downstream copy number or ploidy analyses based on its coverage and heterogeneity level. Therefore, we introduced the homogeneity score (HS) to evaluate the degree of separation between different copy number states (peaks) of the cancer data.

We analyzed low-coverage (< 3x) sequencing data of 20 cancer cell line data with a total of 31 replicates (Supplementary Table 1). We found that HS is weakly correlated with the coverage of the data (Spearman correlation = -0.3104, p-value= 0.0658) (Fig. 2a). Interestingly, we found that HS varies considerably among biological replicates from the same laboratory as well as among the same cell line sequencing data from different studies. For example, biological replicates (denoted with A1 and A2 suffices) from same study exhibit different HS as evidenced from PC3 (78.86%, 71.9%), MCF7 (92.08%, 72.47%), MDA-MB-231 (89.81%, 74.48%), MDA-MB-468 (90.37%, 77.39%) and SK-BR-3 (75.45%, 69.76%). These differences may be partially explained by coverage effect since merging both replicates (denoted with A suffix) slightly improve the HS in the case of PC3 (78.34%), SK-BR-3 (77.54%) and MDA-MBA-468 (92.47%). However, replicate-merging decreases HS in case of MCF7 (66.27%) and MDA-MB-231 (73.07%), suggesting coverage may not be the only key for separating CN profiles into integral CN states. In order to study the relationship between HS and data coverage, we subsampled high-coverage A427 cancer data and found that homogeneity scores remain same for all A427 subsamples (10x to 0.3x) with a maximum change of 2% (Supplementary Table 2). Also, HS is not affected by the whole genome ploidy level (Fig. 2b). Therefore, we concluded that coverage cannot be used as an absolute threshold to assess the suitability of cancer data for copy number analysis. We found that sequencing data with higher HS are less affected by the biases of cancer genomes. After examining all the datasets, we concluded that a minimum homogeneity score of 75% provides an empirical threshold that separates RD frequency distributions peaks and centralized them around CN states. For example, A375_A, A375_B, MDA-MB-468_B, PC3_B, T47D_A, and K562_B data showed low HS and low separation of RD distribution peaks (Supplementary Fig. 1). Out of the 36 samples, aneuploidy profiles of 22 samples with HS > 75 are correctly identified based on the detection of multimodal peaks and known karyotype. The other 14 samples with HS ≤ 75 have 6 accurate and 8 wrong aneuploidy profiles. In such a case, we recommend checking the aneuploidy spectrum manually. Therefore, we argue that homogeneity scoring should be a prerequisite test to be performed on every sequencing data before downstream CNV and aneuploidy analyses.

**Fig. 2.**
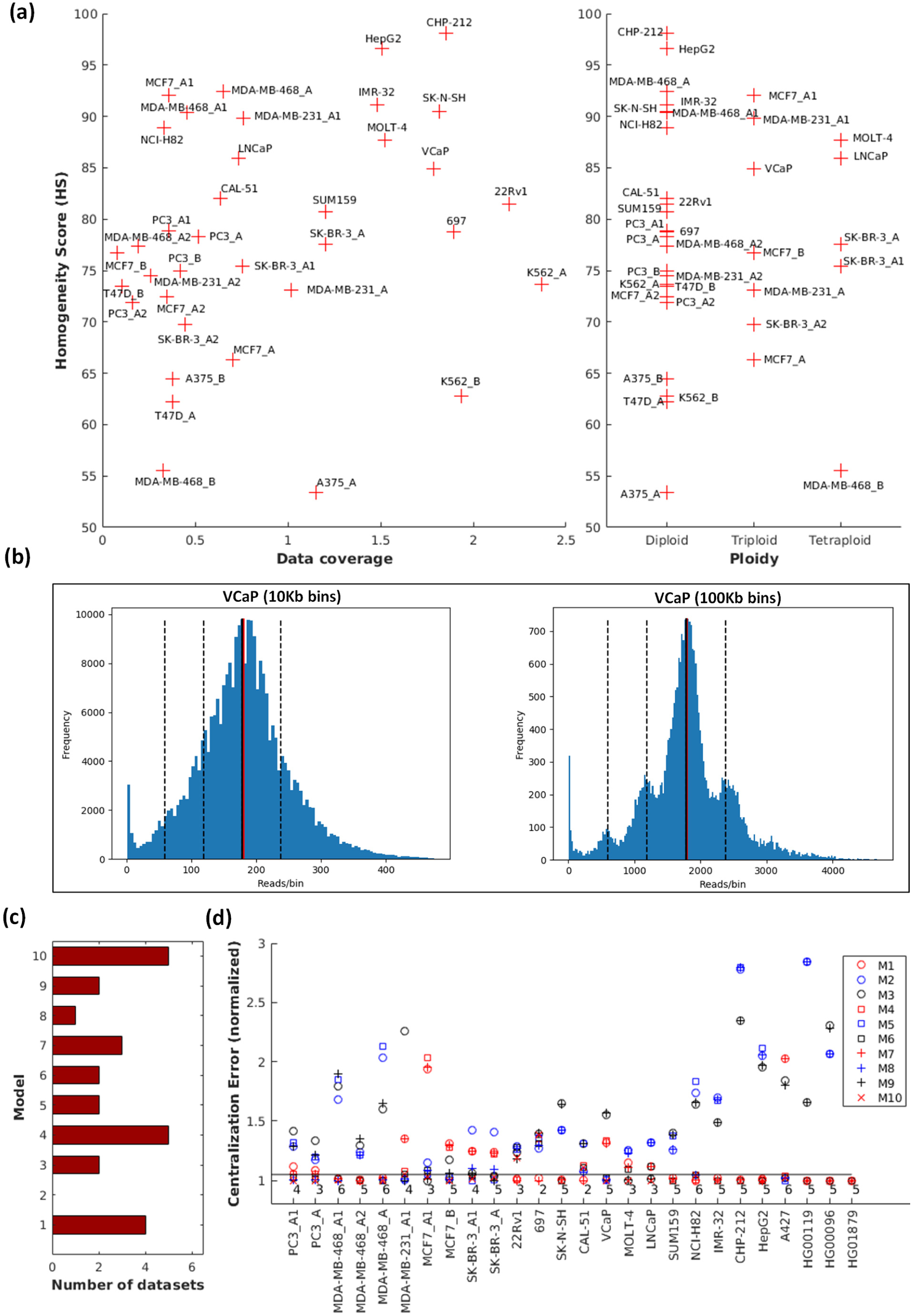
Performance evaluation of AStra. **(a)** Dot plot showing the relationship of the homogeneity score with respect to the data coverage (left) and the ploidy levels (right). **(b)** Frequency distribution of VCaP cancer cell line (left: 10 kb bin; right: 100 kb bin) illustrating AStra can accurately identify the CN states at different resolutions. **(c)** Bar chart showing the number of sequencing datasets (n=26) identified by each model (M1-M10). **(d)** Comparison of centralization errors (CEs) under each model for 26 datasets. The number on the graph denotes the count of models whose CEs lie within 5% difference from the best (lowest) CE.

### 3.2 AStra assumption captures the complex aneuploidy spectrum of cancer cells

Current RD-based ploidy estimation tools compute the whole-genome aneuploidy profile based on best fitting to the empirical models under each ploidy assumptions (diploid, triploid, tetraploid, etc.). However, cancer cells harbor different degrees of hyperploidy that cannot be captured in model with specific ratios of different ploidy states. In contrast, AStra is built on a fundamental assumption that coverage signal of the cancer genome exhibits multimodal distribution with most genomic segments distributed around the integral copy number states. For that, AStra examines many multimodal distributions with different weights of CN states to define the candidate CN reference. CN reference is chosen based on the model that best assimilates the genomic segments (and not RD fitting) into distinct peaks of multimodal distribution. This allows us to apply AStra on low-coverage data and ignore the adverse effects of signal dispersion.

We applied AStra for 36 samples of low-coverage cancer cells, A427 high-coverage cancer cell, and 3 high-coverage normal cells (Supplementary Table 1). AStra computes the aneuploidy profile of low-coverage (<3x) data in less than 3 minutes and high-coverage (28x) in about 15 minutes (Supplementary Table 1). Astra successfully identified the accurate aneuploidy spectrum of all cell lines, with HS greater than 75%, based on two conditions. First, copy number reference and higher CN states are detected as peaks of multimodal distribution (Fig. 2b, Supplementary Fig. 2). Second, the major peak confirms the known karyotype for the ploidy cells (Supplementary Table 1). Similarly, the two major peaks confirm the known karyotype for the hyperploidy cells (Supplementary Table 1). For example, AStra detected the location four peaks (1N, 2N, 3N, 4N) accurately of VCaP at both 10Kb and 100Kb bins (Fig. 2b). Although the RD-frequency of VCaP at 10Kb can be wrongly interpreted as unimodal distribution, AStra still can identify the accurate values of CN states.

We then analyzed the performance of our 10 empirical models for estimating the aneuploidy spectrum by comparing their CEs on different datasets. We also counted the number of datasets that has minimum CE using each model. We found that 9 out of the 10 models results in the minimum CE for at least one dataset (Fig. 2c). M4, M10, and M1 show the best performance as they gave the minimum CE for 5, 5 and 4 datasets, respectively. Remarkably, there is at least one additional model that results in similar CE (<5% difference) for each dataset (Fig. 2d). Therefore, it confirms our hypothesis that aneuploidy spectrum of cancer cells cannot be comprehended based on a single empirical model for each ploidy level. As a conclusion, our models allow AStra to detect the aneuploidy profiles of hyperploidy cancer cells.

### 3.3 AStra best estimates the whole-genome ploidy in low-coverage WGS data

We have also benchmarked our tool against nQuire by applying it on 10 cancer cell lines with known karyotype information and 3 normal cell lines (Supplementary Table 3). Although most cancer cell lines have a hyperploidy profile, nQuire employ models for diploid, triploid, and tetraploid only. Therefore, for a fair comparison, we keep only cancer cell lines which have a major peak at the ploidy CN state and a ratio between genomic segments at major CN state and at any other states is at least 2. This resulted in 6 diploid, 2 triploid and 2 tetraploid cell lines. We found that nQuire estimate the ploidy level of all cells as tetraploid (Supplementary Table 3). In comparison, AStra correctly estimates the nearest whole-genome ploidy level based on the CN state with the highest percentage of genomic segments (Supplementary Table 3).

Although both tools utilize multimodal distribution for estimating the whole-genome aneuploidy, AStra performed better than nQuire. This may be attributed to the fact that Astra evaluates the multimodal fitting based on the genomic segments, not based on the RD signal. This allows Astra to compute the copy number states even at low-coverage data. Also, nQuire assumes unimodal distribution for diploid cells ignoring the existence of any segmental aneuploidy change. Similarly, nQuire triploid distribution assumes that the data distribution can have only two peaks 2N and 3N, and so on. In contrast, AStra models account for any percentage of segmental aneuploidy, and therefore they can capture the complexity of hyperdiploid, near-triploid, or higher ploidy levels.

### 3.4 Aneuploidy spectrum is critical for calibration of CNV detection tools

Most CNV detection tools are built on some basic assumptions or user-defined knowledge of ploidy and percentage of copy number gain/loss for accurate modeling of RD signal data. For example, FREEC uses the whole-genome ploidy level to define the CN state [16] while ReadDepth [15] uses the percentage of gain/loss to adjust their underlying Poisson/negative binomial distribution. Similarly, CNVnator [14] assumes that 99% of the genome is CNV free. However, this information changes across cell lines and between same cell line data obtained from different labs owing to clonal variability as presented in this study (Supplementary Table 1).

The global RD median can be correctly used as ploidy CN state if we assume that the copy number gains and losses are similar. However, this is not the case for most cancer cells. For example, diploid cell lines (CHP-212, IMR-32, HepG2, CAL-51, SUM159) have different percentage of gain (loss) ranging from 1.4% (0%) to 18.11% (3%). Therefore, we compute the median error as the relative shift of the global median (g) to the nearest CN state (s) 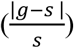. This error can be used to check the validity of ploidy assumption (Fig. 3a). If this error is large (Fig. 3b), the only solution is to correct the copy number of the CNV regions. This can be achieved by computing CN correction factor as the ratio between ploidy state and the global median (Supplementary Table 1). User can use this ratio to adjust the CN of output segments of the tool. Based on these observations, it can be concluded that knowledge of aneuploidy spectrum is fundamental for selecting and calibrating CNV detection tools.

**Fig. 3.**
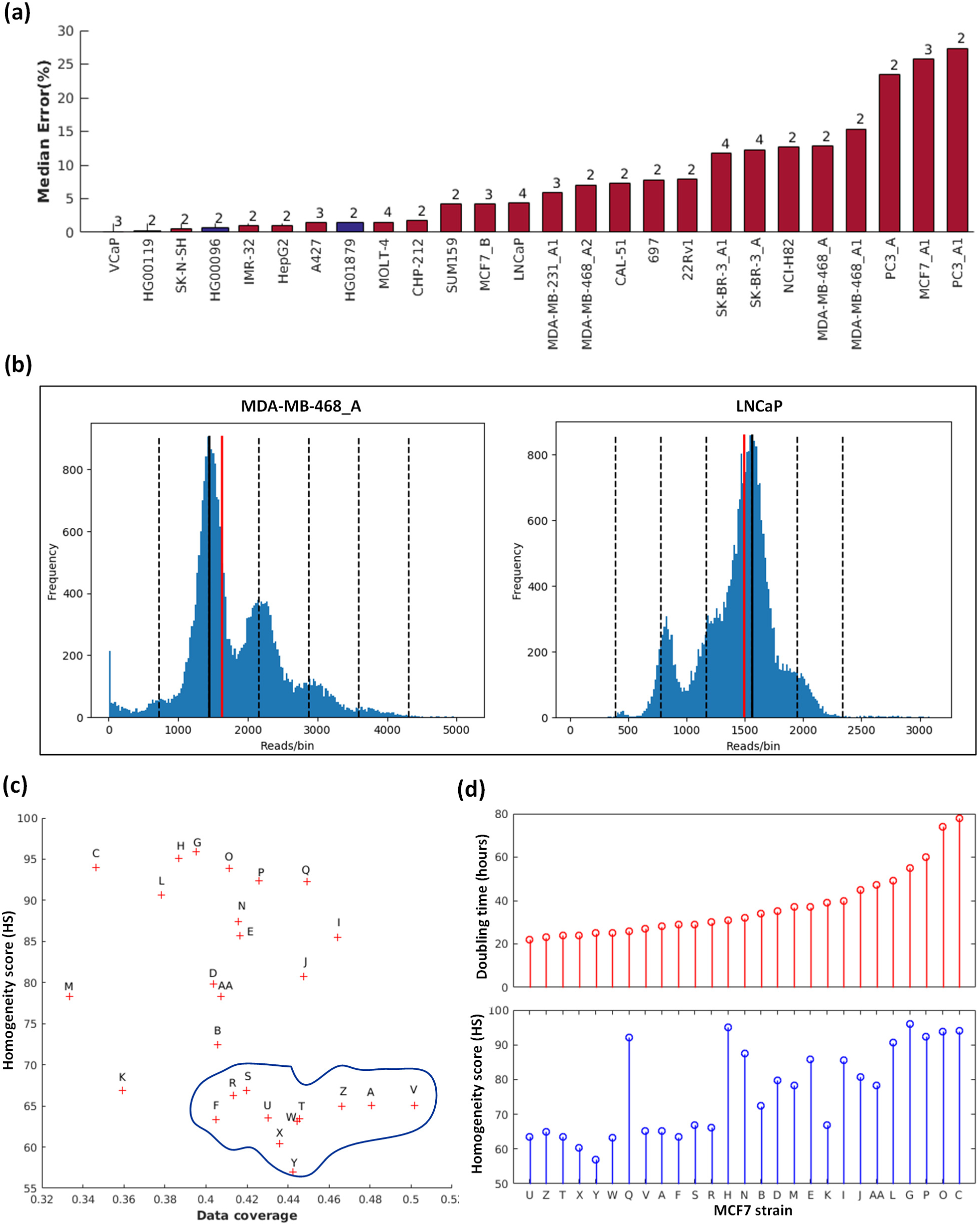
Application of aneuploidy spectrum. **(a)** Bar chart showing the median errors of 26 sequencing datasets of different complexity and ploidy levels. For each dataset, the number on the graph is the estimated ploidy level. The blue bars represent 1000 genomes project samples while red bars represent cancer cell line data. **(b)** Visualization of the median error for MDA-MB-468_A (left) and LNCaP (right) cells. The black line denotes the ploidy state (major CN peak) whereas the red line denotes the global median of RD signal. The dotted black lines denote the other CN states. **(c)** Dot plot showing the relationship between the homogeneity score (HS) and the data coverage for 27 strains of MCF7. The area demarcated by blue line denotes a cluster of strains earlier reported to have similar single nucleotide variant cellular prevalence. **(d)** Comparison of doubling time (top) and the homogeneity score (bottom) of 27 strains of MCF7.

### 3.5 Homogeneity score has biological underpinnings for critical understanding of cancer

Aneuploidy has been linked with cell proliferation defects and lethality in almost all organisms from yeast to human [27, 28]. In mouse and human, chromosomal aneuploidy, with exceptions of few chromosomes, leads to embryonic lethality and the viable trisomies display severe abnormalities [29-31]. Contrasting to its adverse effect on cell proliferation, aneuploidy is associated with uncontrolled cellular growth and cancer [32]. Majority of all human solid tumors carry numerical changes in karyotype [33]. Mouse models of chromosome instability indicate that aneuploidy is directly responsible for cancer formation [34, 35]. In order to understand the effect of aneuploidy profile change on the phenotype of cancer cells, we revisited the study of 27 strains of MCF7 cell lines to obtain the sequencing data [10].

We applied AStra on all the 27 strains of MCF7 collected from different labs (Supplementary Table 4). AStra estimated the aneuploidy spectrum accurately for 26/27 strains with the majority of segments with CN states of 3N or 4N. This confirms that MCF7 karyotype is hypertriploid to hypotetraploid [36]. Out of 27, only the spectrum of strain Z is wrongly identified since it has peaks around the boundaries of CN intervals (2.5N and 3.5N). Interestingly, we also found that strain Z has HS of 65% which is lower than our empirically estimated threshold of >75%. The ploidy levels of the rest 26 strains are inferred as 7 triploid and 19 tetraploid based on the CN state with the maximum percentage of genomic segments (Supplementary Fig. 3). This confirms the idea that populations of the same cell line may have different ploidy profiles based on the culture condition and passage number. Moreover, we found that HS varies greatly among the 26 strains from 56% to 96% with only 14 strains achieving our cutoff threshold of >75% (Fig. 3c). The other 12 strains with low HS suffer from a higher degree of heterogeneity. The RD signal of these heterogeneous strains shows a strong effect of wave artifacts and dispersion (Supplementary Fig. 3d-e). Remarkably, we found that the homogeneity score is correlated with the doubling time (Spearman correlation = 0.7718, p-value = 2.4215e-06) (Fig. 3d) reported earlier for these strains [10]. Additionally, 11 strains (*viz*. A, F, R, S, T, U, V, W, X, Y, and Z) form one group of similar HS and data coverage (Fig. 3c). These same strains were grouped under one cluster in the earlier study based on their SNV (single nucleotide variant) cellular prevalence [10]. This suggests that homogeneity score itself is informative beside the aneuploidy spectrum information.

## 4. Discussion

The genetic profiling of cancer cells is complex. The alterations at the numerical and structural levels and at multitude of length-scales are difficult to detect using a single approach. At the highest level, karyotype alterations include the whole-genome ploidy, chromosomal aneuploidy, and segmental gain/loss. These alterations can limit the accurate estimation of the copy number reference by CNV detection tools. Some CNV detection tools require the prior knowledge of ploidy level while other tools rely on the global median of the coverage signal for estimation of CN reference. The latter can be misleading in most cancer cells. Therefore, we have developed AStra, a Python-based tool, which can segregate the genome into segments of integral CN states. This allows AStra to model the complex hyperploid cancer cells in addition to polyploid cells without any prior knowledge of the input cell line.

Additionally, the heterogeneity of cancer cells adds to another level of complexity. The level of heterogeneity can restrict the legible use of cancer data for aneuploidy profiling. None of the existing ploidy estimation tools provide any quality measurement of the input data. Data coverage is an important factor that affects the quality of the data. However, we showed that coverage cannot be solely relied on as a cut-off threshold. To overcome this problem, AStra defines the homogeneity score (HS) which measures the impact of heterogeneity along with other systematic biases in the cancer data. Our comprehensive analyses of cancer cell lines and visual inspection of the results suggest that a minimum homogeneity score of 75% can be used as a confidence threshold for downstream estimation of ploidy profile.

In conclusion, we recommend using AStra results as a quick reference guide to assess the suitability of cancer NGS data for copy number analysis and to calibrate the parameters of CNV detection tools.

## Declarations

## Acknowledgment

We acknowledge Sanyal and Chattopadhyay lab members for their valuable comments.

## Author’s contributions

AS, AC and AISK conceived the project. AISK developed AStra software with inputs from AS and AC and performed all the analyses. AS, AISK and AC analyzed the data and prepared the manuscript. All authors read and approved the final manuscript.

## Ethics approval and consent to participate

Not applicable.

